# Exploring the effects of BCG vaccination in patients diagnosed with tuberculosis: observational study using the Enhanced Tuberculosis Surveillance system

**DOI:** 10.1101/366476

**Authors:** Sam Abbott, Hannah Christensen, Maeve K Lalor, Dominik Zenner, Colin Campbell, Mary Ramsay, Ellen Brooks-Pollock

**Affiliations:** Bristol Medical School: Population Health Sciences, University of Bristol, Bristol, UK; TB section, National Infection Service, Public Health England, London, UK; Respiratory Diseases Department, Centre for Infectious Disease Surveillance and Control (CIDSC), National Infections Service, Public Health England, London, UK; National Infection Service, Public Health England, London, UK; Immunisation, Hepatitis and Blood safety department, Public Health England, London, UK

**Keywords:** Tuberculosis, BCG, Surveillance, Non-specific, Mortality

## Abstract

**Background:** Bacillus Calmette–Guérin (BCG) is one of the most widely-used vaccines worldwide. BCG primarily reduces the progression from infection to disease, however there is evidence that BCG may provide additional benefits. We aimed to investigate whether there is evidence in routinely-collected surveillance data that BCG vaccination impacts outcomes for tuberculosis (TB) cases in England.

**Methods:** We obtained all TB notifications for 2009–2015 in England from the Enhanced Tuberculosis surveillance system. We considered five outcomes: All-cause mortality, death due to TB (in those who died), recurrent TB, pulmonary disease, and sputum smear status. We used logistic regression, with complete case analysis, to investigate each outcome with BCG vaccination, years since vaccination and age at vaccination, adjusting for potential confounders. All analyses were repeated using multiply imputed data.

**Results:** We found evidence of an association between BCG vaccination and reduced all-cause mortality (aOR:0.76 (95%CI 0.64 to 0.89), P:0.001) and weak evidence of an association with reduced recurrent TB (aOR:0.90 (95%CI 0.81 to 1.00), P:0.056). Analyses using multiple imputation suggested that the benefits of vaccination for all-cause mortality were reduced after 10 years.

**Conclusions:** We found that BCG vaccination was associated with reduced all-cause mortality in people with TB although this benefit was less pronounced more than 10 years after vaccination. There was weak evidence of an association with reduced recurrent TB.

**Highlights:**

- Found evidence of an association between BCG vaccination and reduced all-cause mortality in TB cases.
- Weaker evidence of an association between BCG vaccination and reduced repeat TB episodes in TB cases.
- There was little evidence of an association with other TB outcomes.
- We explored the identified associations by age and time since vaccination.

## INTRODUCTION

Bacillus Calmette–Guérin (BCG) is one of the mostly widely-used vaccines and the only vaccine that protects against tuberculosis (TB) disease. BCG was first used in humans in 1921 and was introduced into the WHO Expanded Program on Immunization in 1974.[1] BCG vaccination has been controversial due to its variable efficacy and possibility of causing a false positive result with the standard skin test for TB.[2] However, the lack of a more effective vaccine and the emergence of drug-resistant TB strains means that BCG remains the best available vaccination for TB.

BCG’s primary mode of action is to directly prevent the development of active, symptomatic disease. Its efficacy in adults is context specific, with estimates ranging between 0% and 78%.[3] Efficacy has been shown to be dependent on previous exposure, with unexposed individuals receiving the greatest benefit.[4] Unlike in adults, BCG has consistently been shown to be highly protective against TB and TB meningitis in children.[5,6] For this reason the majority of countries that use BCG vaccinate at birth.[7,8] Adult vaccination is no longer common in the UK, where universal BCG vaccination of adolescents was stopped in 2005 in favour of a targeted neonatal programme aimed at high risk children.

Vaccination policy has been primarily based on reducing the incidence of active TB and little attention has been given to any additional effects of BCG.[9,10] There is some evidence that BCG vaccination induces innate immune responses which may provide non-specific protection,[11] TB patients with BCG scars were found to respond better to treatment with earlier sputum smear conversion,[12] and there is evidence to suggest that BCG vaccination is associated with reduced all-cause neonatal mortality[13,14] and both reduced TB[15] and all-cause[16] mortality in the general population. Given that the immunology behind TB immunity is not well understood these findings suggest that BCG may play a more important role in improving TB outcomes than previously thought. We aimed to quantify the effects of BCG vaccination on outcomes for individuals with notified TB in England using routinely collected surveillance data to provide evidence for appropriate public health action and provision. Where we found an association, we additionally explored the role of years since vaccination, and age at vaccination.

## METHOD

### Enhanced Tuberculosis Surveillance (ETS) system

We extracted all notifications from the Enhanced Tuberculosis Surveillance (ETS) system from January 1, 2009 to December 31, 2015. BCG vaccination status and year of vaccination have been collected since 2008. We considered five TB outcomes which were selected due to their association with increased case infectiousness or poor consequences for patients (table 1).

**Table 1:** Summary of outcome definitions and rationale for inclusion

| Outcome | Definition |
| --- | --- |
| All-cause mortality | All-cause mortality was defined using the overall outcome recorded in ETS, this is based on follow up at 12, 24, and 36 months (or until treatment completion). If a case dies within 12 months of completing treatment, from TB or from a cause related to TB then their overall outcome will be updated in the ETS. Those that were lost to follow up, or not evaluated were treated as missing. |
| Death due to TB (in those who died) | For cases with a known cause of death, death due to TB is defined as those that died from TB, or where TB had contributed to their death. |
| Recurrent TB | TB cases who had recurrent episodes were identified in the dataset. |
| Pulmonary disease | Cases that present with pulmonary TB (regardless of extra-pulmonary TB status). |
| Sputum smear status - positive | Status of sputum sample tested for Acid-Fast Bacilli. |

### Exposure variables relating to BCG

We included three exposure variables related to BCG: BCG status (vaccinated, yes/no), years since vaccination and age at vaccination.

BCG status was taken directly from the ETS. Years since BCG vaccination was defined as year of notification minus year of vaccination and categorised into two groups (0 to 10 and 11+ years), based on evidence that the average duration of BCG protection is 10–15 years.[15] Age at vaccination is defined in the online supplementary information.

### Statistical Analysis

R was used for all statistical analysis.[17] The analysis was conducted in two stages. Firstly, we calculated proportions for all demographic and outcome variables, and compared vaccinated and unvaccinated TB cases using the *χ*^2^ test. Secondly, we used logistic regression, with complete case analysis, to estimate the association between exposures and outcome variables, both with and without adjustment for confounders.

In the multivariable models, we adjusted for sex,[18–20] age,[21] Index of Multiple Deprivation (2010) categorised into five groups for England (IMD rank),[22,23] ethnicity,[18,24] UK birth status,[25,26] and year of notification. As the relationship between age and outcomes was non-linear, we modelled age using a natural cubic spline with knots at the 25%, 50% and 75% quantiles.

We conducted sensitivity analyses to assess the robustness of the results, by dropping each confounding variable in turn and assessing the effect on the adjusted Odds Ratios (aORs) of the exposure variable. We repeated the analysis excluding duplicate recurrent cases, and restricting the study population to those eligible for the BCG schools scheme (defined as UK born cases that were aged 14 or over in 2004) to assess the comparability of the BCG vaccinated and unvaccinated populations. To mitigate the impact of missing data we used multiple imputation, with the MICE package.[27] We imputed 50 data sets (for 20 iterations) using all variables included in the analysis as predictors along with Public Health England centre. The model results were pooled using the small sample method,[28] and effect sizes compared with those from the main analysis.

## RESULTS

### Description of the data

There were 51,645 TB notifications between 2009–2015 in England. Reporting of vaccination status and year of vaccination improved over time: 64.9% (20865/32154) of notifications included vaccination status for 2009 to 2012, increasing to 70% (13647/19491) from 2013 to 2015. The majority of cases that had a known vaccination status were vaccinated (70.6%, 24354/34512), and where age and year of vaccination was known, the majority of cases were vaccinated at birth (60%, 5979/10066).

Vaccinated cases were younger than unvaccinated cases on average (median age 34 years (IQR 26 to 45) compared to 38 years (IQR 26 to 62)). A higher proportion of non-UK born cases were BCG vaccinated, (72.7%, 18297/25171) compared to UK born cases (65.2%, 5787/8871, P: < 0.001) and, of those vaccinated, a higher proportion of non-UK born cases were vaccinated at birth compared to UK born cases (68%, 4691/6896 vs. 40.5%, 1253/3096 respectively, P: < 0.001). See online supplementary table S1 for the breakdown of outcome variables and supplementary table S2 for the breakdown of confounding variables.

### All-cause mortality

In the univariable analysis the odds of death from any cause were lower for BCG vaccinated TB cases compared to unvaccinated cases, with an OR of 0.28 (95% CI 0.24 to 0.32, P: <0.001) (table 2, see supplementary table S3 for the full table); an association remained after adjusting for confounders, but was attenuated with an aOR of 0.76 (95% CI 0.64 to 0.89, P: 0.001). We estimate that if all unvaccinated cases had been vaccinated there would have been on average 19 (95% CI 9 to 29) fewer deaths per year during the study period (out of 81 deaths per year on average in unvaccinated cases). Whilst there was evidence in univariable analyses to suggest all-cause mortality was higher in persons vaccinated more than 10 years prior to notification of TB and that all-cause mortality increased with increasing age group, these disappeared after adjusting for potential confounders (table 3, supplementary table S4).

Similar results to the multivariable analysis were found using multiply imputed data for the association between vaccination status and all-cause mortality (aOR: 0.76 (95% CI 0.61 to 0.94), P: 0.013), but not for time since vaccination with a greatly increased risk of all-cause mortality estimated for those vaccinated more than 10 years before case notification, compared to those vaccinated more recently (aOR: 12.19 (95% CI 3.48 to 42.64), (see online supplementary table S5, supplementary table S6)). For age at vaccination results for the multivariable analysis using multiply imputed data were comparable to those found using complete case analysis, except that there was some evidence that vaccination in adolescence, compared to under 1, was associated with increased, rather than decreased, all-cause mortality (aOR: 1.57 (95% CI 1.13 to 2.19), supplementary table S7).

### Deaths due to TB (in those who died)

There was little evidence of any association between BCG vaccination and deaths due to TB (in those who died and where cause of death was known) in the univariable analysis (table 2). The adjusted point estimate indicated an association between BCG vaccination and reduced deaths due to TB (in those who died) although the confidence intervals remained wide with a similar result found using multiply imputed data (see online supplementary table S5). There were insufficient data to robustly estimate an association between deaths due to TB (in those who died) and years since vaccination or age at vaccination (table 3, supplementary table S4).

### Recurrent TB

In both the univariable and multivariable analysis there was some evidence that BCG vaccination was associated with reduced recurrent TB, although the strength of the evidence was weakened after adjusting for confounders (table 2). In the adjusted analysis, the odds of recurrent TB were lower for BCG vaccinated cases compared to unvaccinated cases, with an aOR of 0.90 (95% CI 0.81 to 1.00, P: 0.056). The strength of the evidence for this association was comparable in the analysis using multiply imputed data (see online supplementary table S5). There was little evidence in the adjusted analysis of any association between recurrent TB and years since vaccination (table 3) or age at vaccination (supplementary table S4).

### Other Outcomes

After adjusting for confounders there was little evidence for any association between BCG vaccination and pulmonary disease or positive sputum smear status (table 2); similar results were found using multiply imputed data (see online supplementary table S5).

**Table 2:**
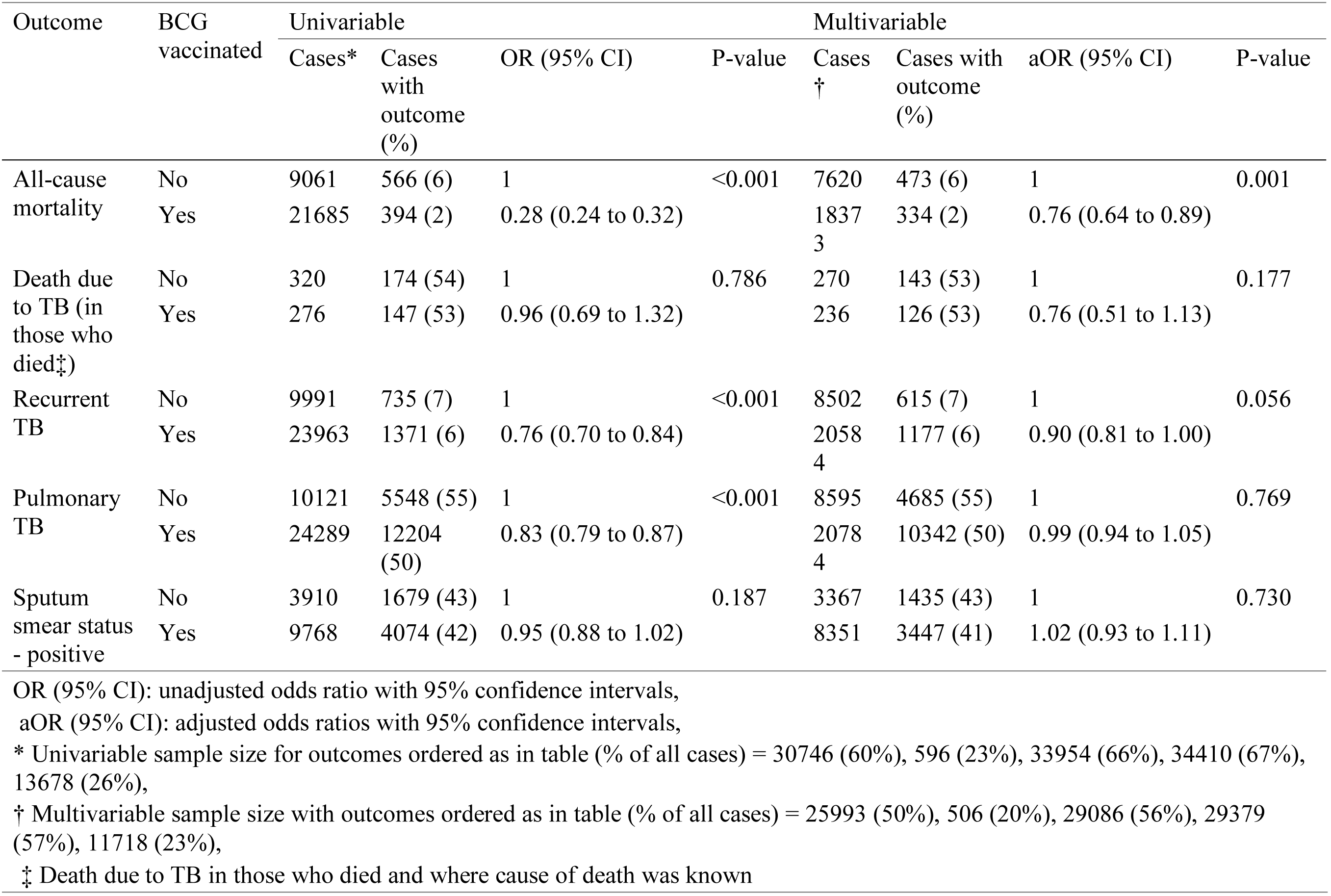
Summary of associations between BCG vaccination and all outcomes

| Outcome | BCG vaccinated | Univariable |  |  |  | Multivariable |  |  |  |
| --- | --- | --- | --- | --- | --- | --- | --- | --- | --- |
|  |  | Cases* | Cases with outcome (%) | OR (95% CI) | P-value | Cases † | Cases with outcome (%) | aOR (95% CI) | P-value |
| All-cause mortality | No | 9061 | 566 (6) | 1 | <0.001 | 7620 | 473 (6) | 1 | 0.001 |
|  | Yes | 21685 | 394 (2) | 0.28 (0.24 to 0.32) |  | 1837<br>3 | 334 (2) | 0.76 (0.64 to 0.89) |  |
| Death due to TB (in those who died‡) | No | 320 | 174 (54) | 1 | 0.786 | 270 | 143 (53) | 1 | 0.177 |
|  | Yes | 276 | 147 (53) | 0.96 (0.69 to 1.32) |  | 236 | 126 (53) | 0.76 (0.51 to 1.13) |  |
| Recurrent TB | No | 9991 | 735 (7) | 1 | <0.001 | 8502 | 615 (7) | 1 | 0.056 |
|  | Yes | 23963 | 1371 (6) | 0.76 (0.70 to 0.84) |  | 2058<br>4 | 1177 (6) | 0.90 (0.81 to 1.00) |  |
| Pulmonary TB | No | 10121 | 5548 (55) | 1 | <0.001 | 8595 | 4685 (55) | 1 | 0.769 |
|  | Yes | 24289 | 12204 (50) | 0.83 (0.79 to 0.87) |  | 2078<br>4 | 10342 (50) | 0.99 (0.94 to 1.05) |  |
| Sputum smear status - positive | No | 3910 | 1679 (43) | 1 | 0.187 | 3367 | 1435 (43) | 1 | 0.730 |
|  | Yes | 9768 | 4074 (42) | 0.95 (0.88 to 1.02) |  | 8351 | 3447 (41) | 1.02 (0.93 to 1.11) |  |

OR (95% CI): unadjusted odds ratio with 95% confidence intervals,
aOR (95% CI): adjusted odds ratios with 95% confidence intervals,
\* Univariable sample size for outcomes ordered as in table (% of all cases) = 30746 (60%), 596 (23%), 33954 (66%), 34410 (67%), 13678 (26%),
† Multivariable sample size with outcomes ordered as in table (% of all cases) = 25993 (50%), 506 (20%), 29086 (56%), 29379 (57%), 11718 (23%),
‡ Death due to TB in those who died and where cause of death was known

**Table 3:** Summary of associations between years since vaccination and all outcomes in individuals who were vaccinated. The baseline exposure is vaccination ≤ 10 years before diagnosis compared to vaccination 11 + years before diagnosis. Deaths due to TB (in those who died) had insufficient data for effect sizes to be estimated in both the univariable and multivariable analysis.

| Outcome | Years since BCG | Univariable |  |  |  | Multivariable |  |  |  |
| --- | --- | --- | --- | --- | --- | --- | --- | --- | --- |
|  |  | Cases* | Cases with outcome (%) | OR (95% CI) | P-value | Cases† | Cases with outcome (%) | aOR (95% CI) | P-value |
| All-cause mortality | $\leq 10$ | 718 | 5 (1) | 1 | 0.004 | 554 | 4 (1) | 1 | 0.897 |
|  | 11+ | 8106 | 166 (2) | 2.98 (1.22 to 7.28) |  | 7171 | 148 (2) | 0.91 (0.24 to 3.54) |  |
| Death due to TB (in those who died‡) | $\leq 10$ | 2 | 2 (100) | 1 | - | 2 | 2 (100) | 1 | - |
|  | 11+ | 108 | 59 (55) | *Insufficient data* |  | 98 | 53 (54) | *Insufficient data* |  |
| Recurrent TB | $\leq 10$ | 780 | 22 (3) | 1 | 0.005 | 613 | 14 (2) | 1 | 0.515 |
|  | 11+ | 9172 | 451 (5) | 1.78 (1.15 to 2.75) |  | 8194 | 406 (5) | 1.24 (0.63 to 2.44) |  |
| Pulmonary TB | $\leq 10$ | 770 | 480 (62) | 1 | <0.001 | 601 | 382 (64) | 1 | 0.309 |
|  | 11+ | 9248 | 4757 (51) | 0.64 (0.55 to 0.74) |  | 8254 | 4232 (51) | 0.87 (0.67 to 1.14) |  |
| Sputum smear status - positive | $\leq 10$ | 157 | 81 (52) | 1 | 0.941 | 122 | 61 (50) | 1 | 0.920 |
|  | 11+ | 3064 | 1590 (52) | 1.01 (0.73 to 1.40) |  | 2734 | 1405 (51) | 1.02 (0.68 to 1.54) |  |

OR (95% CI): unadjusted odds ratio with 95% confidence intervals, aOR (95% CI): adjusted odds ratios with 95% confidence intervals,
\* Univariable sample size for outcomes ordered as in table (% of vaccinated cases) = 8824 (36%), 110 (28%), 9952 (41%), 10018 (41%), 3221 (13%),
† Multivariable sample size with outcomes ordered as in table (% of vaccinated cases) = 7725 (32%), 100 (25%), 8807 (36%), 8855 (36%), 2856 (12%),
‡ Death due to TB in those who died and where cause of death was known

### Sensitivity analysis

Dropping duplicate recurrent TB notifications increased the magnitude, and precision, of the effect sizes for recurrent TB, all-cause mortality, and deaths due to TB (in those who died) (see online supplementary table S8). Restricting the analysis to only cases that were eligible for the BCG schools scheme reduced the sample size of the analysis (from an initial study size of 51645, of which 12832 were UK born, to 9943 cases that would have been eligible for the BCG schools scheme). With this reduced sample size, there was strong evidence in adjusted analyses of an association between BCG vaccination and reduced recurrent TB, and evidence of an association with decreased all-cause mortality (see online supplementary table S8).

## DISCUSSION

Using TB surveillance data collected in England we found that BCG vaccination, prior to the development of active TB, was associated with reduced all-cause mortality and fewer recurrent TB cases, although the evidence for this association was weaker. There was some suggestion that the association with all-cause mortality was due to reduced deaths due to TB (in those who died), though the study was underpowered to definitively assess this. We did not find evidence of an association between BCG status and positive smear status or pulmonary TB. Analysis with multiply imputed data indicated that notification 10+ years after vaccination was associated with increased all-cause mortality. In separate analyses, there was some evidence that vaccination at birth, compared to at any other age, was associated with reduced all-cause mortality, and increased deaths due to TB (in those who died).

This study used a large detailed dataset, with coverage across demographic groups, and standardized data collection from notifications and laboratories. The use of routine surveillance data means that this study would be readily repeatable with new data. The surveillance data contained multiple known risk factors, this allowed us to adjust for these confounders in the multivariable analysis, which attenuated the evidence for an association with BCG vaccination for all outcomes. However, there are important limitations to consider. The study was conducted within a population of active TB cases, therefore the association with all-cause mortality cannot be extrapolated to the general population. Additionally, vaccinated and unvaccinated populations may not be directly comparable because vaccination has been targeted at high-risk neonates in the UK since 2005. We mitigated this potential source for bias by conducting a sensitivity analysis including only those eligible for the universal school age scheme, and whilst the strength of associations were attenuated there remained some evidence of improved outcomes. Sensitivity analysis excluding recurrent cases indicated their inclusion may have biased our results towards the null.

Variable data completeness changed with time, with both BCG vaccination status and year of vaccination having a high percentage of missing data, which may not be missing completely at random. We therefore checked the robustness of our results with multiple imputation including regional variability, however an unknown missing not at random mechanism, or unmeasured confounding may still have introduced bias. We found a greatly increased risk of all-cause mortality for those vaccinated more than 10 years ago in the analysis with multiply imputed data, compared to the complete case analysis. This is likely to be driven by a missing not at random mechanism for years since vaccination, with older cases being both more likely to have been vaccinated more than 10 years previously and to also have an unknown year of vaccination. The high percentage of missing data also means that we were likely to be underpowered to detect an effect of BCG vaccination on sputum smear status and deaths due to TB (in those who died), with years since vaccination, and age at vaccination likely to be underpowered for all outcomes. We were not able to adjust for either tuberculin skin test (TST) stringency, or the latitude effect, although we were able to adjust for UK birth status.[29] However, the bias induced by these confounders is likely to be towards the null, meaning that our effect estimates are likely to be conservative. Finally, BCG vaccination status may be subject to misclassification due to recall bias; validation studies of the recording of BCG status in the ETS would be required to assess this.

Little work has been done to assess the overall effect of BCG on outcomes for active TB cases although the possible non-specific effects of BCG are an area of active research.[14,30,31] Whilst multiple studies have investigated BCG’s association with all-cause mortality, it has been difficult to assess whether the association continues beyond the first year of life.[31] The effect size of the association we identified between BCG and all-cause mortality in active TB cases was comparable to that found in a Danish case-cohort study in the general population (aHR: 0.58 (95% CI 0.39 to 0.85).[16] A recent systematic review also found that BCG vaccination was associated with reduced all-cause mortality in neonates, with an average relative risk of 0.70 (95% CI 0.49 to 1.01) from five clinical trials and 0.47 (95% CI 0.32 to 0.69) from nine observational studies at high risk of bias.[14] We found some weak evidence that BCG vaccination was associated with reduced deaths due to TB (in those who died), although our point estimate had large confidence intervals. Several meta-analyses have found evidence supporting this association,[6,15] with one meta-analysis estimating a 71% (RR: 0.29 95% CI 0.16 to 0.53) reduction in deaths due to TB in individuals vaccinated with BCG.[6] The meta-analysis performed by Abubakar et al. also found consistent evidence for this association, with a Rate Ratio of 0.22 (95% CI 0.15 to 0.33).[15] In contrast to our study, both of these meta-analyses estimated the protection from TB mortality in BCG vaccinated individuals rather than in BCG vaccinated cases who had died from any cause. Additionally, neither study explored the association between BCG vaccination and all-cause mortality or recurrent TB. This study could not determine the possible causal pathway for the association between BCG vaccination all-cause mortality, and recurrent TB. These are important to establish in order to understand the effect of BCG vaccination on TB outcomes.

We found that BCG vaccination was associated with reduced all-cause mortality, with some weaker evidence of an association with reduced recurrent TB. A plausible mechanism for this association is that BCG vaccination improves treatment outcomes,[12] which then results in decreased mortality, and reduced recurrent TB. However, these effects may also be independent and for all-cause mortality may not be directly related to active TB. In this case, a possible mechanism for the association between BCG vaccination and all-cause mortality is that BCG vaccination modulates the innate immune response, resulting in non-specific protection.[11] For low incidence countries, where the reduction in TB cases has been used as evidence to scale back vaccination programs,[7] these results suggest that BCG vaccination may be more beneficial than previously thought. In countries that target vaccination at those considered to be at high risk of TB the results from this study could be used to help drive uptake by providing additional incentives for vaccination. The evidence we have presented should be considered in future cost-effectiveness studies of BCG vaccination programs.

Further work is required to determine whether years since vaccination and age at vaccination are associated with TB outcomes as this study was limited by low sample size, missing data for year of vaccination, and the relative rarity of some TB outcomes. However, due to the continuous collection of the surveillance data used in this analysis, this study could be repeated once additional data have been collected. The results from this study require validation in independent datasets and the analysis should be reproducible in other low incidence countries that have similarly developed surveillance systems. If validated in low incidence countries, similar studies in medium to high incidence countries should be conducted because any effect would have a greater impact in these settings.

## Acknowledgements

The authors thank the TB section at Public Health England (PHE) for maintaining the Enhanced Tuberculosis Surveillance (ETS) system; all the healthcare workers involved in data collection for the ETS.

## Contributors

SA, HC, and EBP conceived and designed the work. SA undertook the analysis with advice from all other authors. All authors contributed to the interpretation of the data. SA wrote the first draft of the paper and all authors contributed to subsequent drafts. All authors approve the work for publication and agree to be accountable for the work.

## Accessibility of data and programming code

The code for the analysis contained in this paper can be found at: doi.org/10.5281/zenodo.1213799

## Online supplementary appendix: Exploring the effects of BCG vaccination in patients diagnosed with tuberculosis: observational study using the Enhanced Tuberculosis Surveillance system

### Outcome variables stratified by BCG vaccination status

**Supplementary table S1:** Outcomes for individuals in England notified with tuberculosis between 2009–2015, stratified by BCG vaccination status.

| Outcome | Total | Vaccinated | Unvaccinated | Unknown vaccine status |
| --- | --- | --- | --- | --- |
| Total, all cases | 51645 | 24354 {47} | 10158 {20} | 17133 {33} |
| All-cause mortality | 45588 (88) | 21685 (89) | 9061 (89) | 14842 (87) |
| No | 43024 [94] | 21291 [98] | 8495 [94] | 13238 [89] |
| Yes | 2564 [6] | 394 [2] | 566 [6] | 1604 [11] |
| Death due to TB (in those who died*) | 1373 (3) | 276 (1) | 320 (3) | 777 (5) |
| No | 572 [42] | 129 [47] | 146 [46] | 297 [38] |
| Yes | 801 [58] | 147 [53] | 174 [54] | 480 [62] |
| Recurrent TB | 48497 (94) | 23963 (98) | 9991 (98) | 14543 (85) |
| No | 44869 [93] | 22592 [94] | 9256 [93] | 13021 [90] |
| Yes | 3628 [7] | 1371 [6] | 735 [7] | 1522 [10] |
| Pulmonary TB | 51432 (100) | 24289 (100) | 10121 (100) | 17022 (99) |
| Extra-pulmonary (EP) only | 24280 [47] | 12085 [50] | 4573 [45] | 7622 [45] |
| Pulmonary, with or without EP | 27152 [53] | 12204 [50] | 5548 [55] | 9400 [55] |
| Sputum smear status - positive | 19551 (38) | 9768 (40) | 3910 (38) | 5873 (34) |
| Negative | 11060 [57] | 5694 [58] | 2231 [57] | 3135 [53] |
| Positive | 8491 [43] | 4074 [42] | 1679 [43] | 2738 [47] |

{% all cases}{% complete within vaccine status}[% complete within category],
\* Death due to TB in those who died and where cause of death was known

### Confounding variables stratified by BCG vaccination status

**Supplementary table S2:** Confounders for individuals in England notified with tuberculosis between 2009–2015, stratified by BCG vaccination status.

| Confounder | Total | Vaccinated | Unvaccinated | Unknown vaccine status |
| --- | --- | --- | --- | --- |
| Total, all cases | 51645 | 24354 {47} | 10158 {20} | 17133 {33} |
| Age | 51645 (100) | 24354 (100) | 10158 (100) | 17133 (100) |
| Mean [SD] | 40 [19] | 36 [16] | 44 [22] | 45 [20] |
| Median [25%, 75%] | 36 [27, 52] | 34 [26, 45] | 38 [26, 62] | 41 [29, 59] |
| Sex | 51535 (100) | 24320 (100) | 10136 (100) | 17079 (100) |
| Female | 22066 [43] | 10791 [44] | 4312 [43] | 6963 [41] |
| Male | 29469 [57] | 13529 [56] | 5824 [57] | 10116 [59] |
| IMD rank (with 1 as most deprived and 5 as least deprived) | 43525 (84) | 21240 (87) | 8866 (87) | 13419 (78) |
| 1 | 16800 [39] | 7779 [37] | 3665 [41] | 5356 [40] |
| 2 | 13057 [30] | 6836 [32] | 2564 [29] | 3657 [27] |
| 3 | 6838 [16] | 3459 [16] | 1259 [14] | 2120 [16] |
| 4 | 4045 [9] | 1893 [9] | 836 [9] | 1316 [10] |
| 5 | 2785 [6] | 1273 [6] | 542 [6] | 970 [7] |
| UK born | 49820 (96) | 24084 (99) | 9958 (98) | 15778 (92) |
| Non-UK Born | 36988 [74] | 18297 [76] | 6874 [69] | 11817 [75] |
| UK Born | 12832 [26] | 5787 [24] | 3084 [31] | 3961 [25] |
| Ethnic group | 50416 (98) | 24074 (99) | 10024 (99) | 16318 (95) |
| White | 10194 [20] | 3560 [15] | 2695 [27] | 3939 [24] |
| Black-Caribbean | 1112 [2] | 559 [2] | 242 [2] | 311 [2] |
| Black-African | 8942 [18] | 4620 [19] | 1602 [16] | 2720 [17] |
| Black-Other | 462 [1] | 261 [1] | 80 [1] | 121 [1] |
| Indian | 12994 [26] | 7176 [30] | 2061 [21] | 3757 [23] |
| Pakistani | 8237 [16] | 3512 [15] | 1720 [17] | 3005 [18] |
| Bangladeshi | 2025 [4] | 918 [4] | 480 [5] | 627 [4] |
| Chinese | 601 [1] | 289 [1] | 101 [1] | 211 [1] |
| Mixed / Other | 5849 [12] | 3179 [13] | 1043 [10] | 1627 [10] |
| Calendar year | 51645 (100) | 24354 (100) | 10158 (100) | 17133 (100) |

{% all cases}{% complete within vaccine status}[% complete within category],
\* Death due to TB in those who died and where cause of death was known

### Model output: All-cause mortality

**Supplementary table S3:**
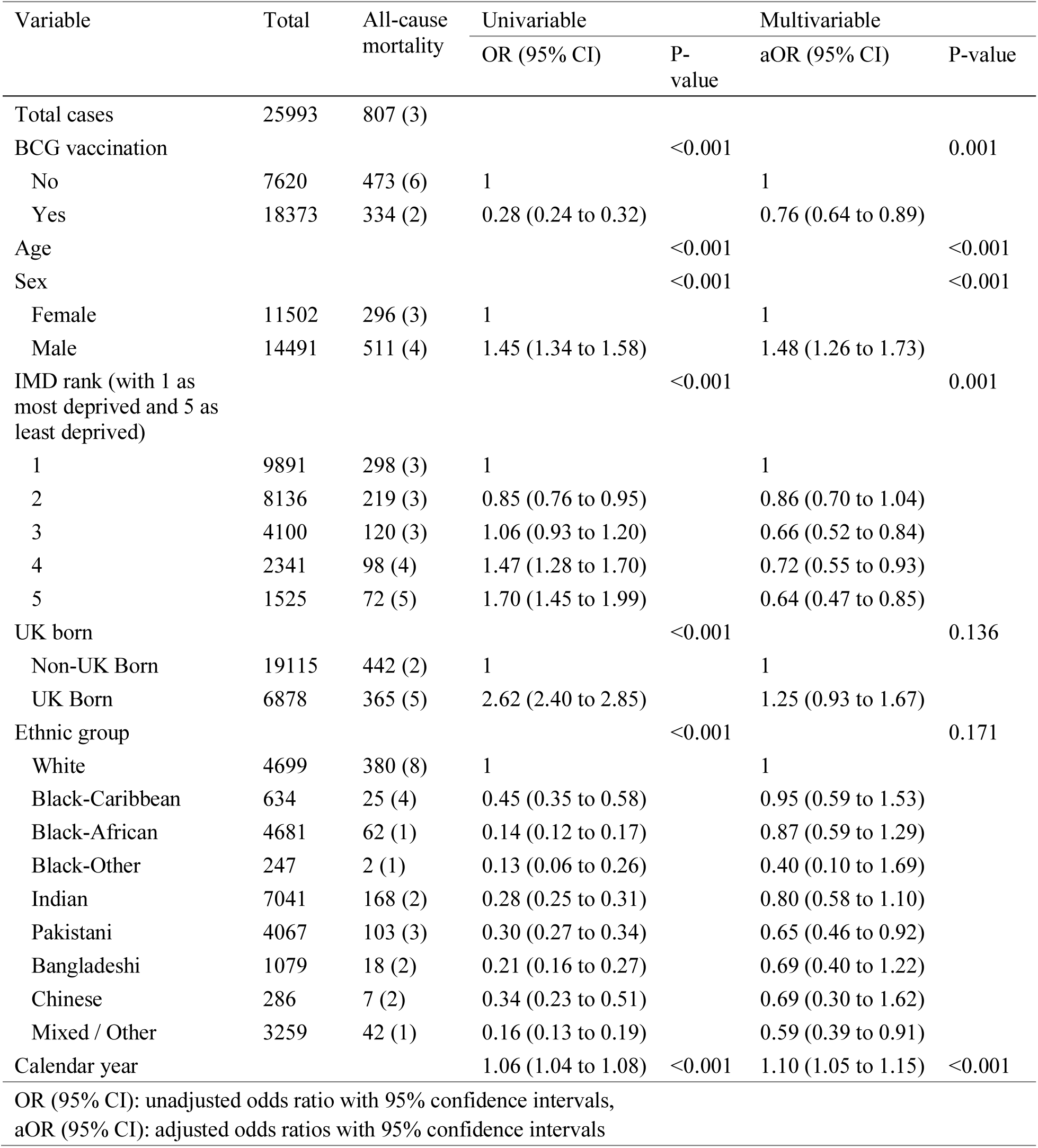
Summary of logistic regression model output with BCG vaccination as the exposure and all-cause mortality as the outcome.

### Secondary exposure: Age at vaccination

We calculated age at vaccination as year of vaccination minus year of birth. We categorized age at vaccination into 0 to < 1, 1 to < 12, 12 to < 16 and 16+ years because the distribution was bi-model with modes at 0 and 12 years. This categorization captures the current UK policy of vaccination at birth, historic policy of vaccination at 13–15 years and catch up vaccination for high risk children.

**Supplementary table S4:** Summary of associations between age at vaccination and all outcomes in individuals who were vaccinated - the baseline exposure is vaccination at birth compared to vaccination from 1 to < 12, 12 to < 16, and 16+ years of age.

| Outcome | Age at BCG | Univariable |  |  |  | Multivariable |  |  |  |
| --- | --- | --- | --- | --- | --- | --- | --- | --- | --- |
|  |  | Cases* | Cases with outcome (%) | OR (95% CI) | P-value | Cases† | Cases with outcome (%) | aOR (95% CI) | P-value |
| All-cause mortality | < 1 | 5234 | 45 (1) | 1 | <0.001 | 4626 | 43 (1) | 1 | 0.127 |
|  | 1 to < 12 | 1915 | 58 (3) | 3.60 (2.43 to 5.34) |  | 1678 | 52 (3) | 1.36 (0.85 to 2.16) |  |
|  | 12 to < 16 | 1267 | 41 (3) | 3.86 (2.51 to 5.91) |  | 1094 | 32 (3) | 0.81 (0.45 to 1.46) |  |
|  | ≥ 16 | 408 | 27 (7) | 8.17 (5.01 to 13.32) |  | 327 | 25 (8) | 1.41 (0.76 to 2.63) |  |
| Death due to TB (in those who died‡) | < 1 | 27 | 20 (74) | 1 | 0.118 | 27 | 20 (74) | 1 | 0.543 |
|  | 1 to < 12 | 43 | 20 (47) | 0.30 (0.11 to 0.87) |  | 39 | 18 (46) | 0.36 (0.08 to 1.51) |  |
|  | 12 to < 16 | 23 | 13 (57) | 0.46 (0.14 to 1.50) |  | 17 | 9 (53) | 0.40 (0.06 to 2.52) |  |
|  | ≥ 16 | 17 | 8 (47) | 0.31 (0.09 to 1.12) |  | 17 | 8 (47) | 0.35 (0.06 to 2.16) |  |
| Recurrent TB | < 1 | 5909 | 284 (5) | 1 | 0.463 | 5275 | 258 (5) | 1 | 0.246 |
|  | 1 to < 12 | 2174 | 105 (5) | 1.01 (0.80 to 1.26) |  | 1928 | 92 (5) | 0.84 (0.65 to 1.09) |  |
|  | 12 to < 16 | 1421 | 58 (4) | 0.84 (0.63 to 1.12) |  | 1242 | 51 (4) | 0.70 (0.48 to 1.02) |  |
|  | ≥ 16 | 448 | 26 (6) | 1.22 (0.81 to 1.85) |  | 362 | 19 (5) | 0.82 (0.49 to 1.37) |  |
| Pulmonary TB | < 1 | 5946 | 2828 (48) | 1 | <0.001 | 5305 | 2510 (47) | 1 | 0.005 |
|  | 1 to < 12 | 2194 | 1159 (53) | 1.23 (1.12 to 1.36) |  | 1941 | 1033 (53) | 1.15 (1.02 to 1.29) |  |
|  | 12 to < 16 | 1425 | 971 (68) | 2.36 (2.09 to 2.67) |  | 1245 | 846 (68) | 1.09 (0.92 to 1.29) |  |
|  | ≥ 16 | 453 | 279 (62) | 1.77 (1.45 to 2.15) |  | 364 | 225 (62) | 1.47 (1.15 to 1.88) |  |
| Sputum smear status - positive | < 1 | 1753 | 836 (48) | 1 | <0.001 | 1557 | 742 (48) | 1 | 0.862 |
|  | 1 to < 12 | 755 | 394 (52) | 1.20 (1.01 to 1.42) |  | 682 | 348 (51) | 0.96 (0.79 to 1.17) |  |
|  | 12 to < 16 | 556 | 357 (64) | 1.97 (1.62 to 2.40) |  | 486 | 308 (63) | 1.06 (0.81 to 1.39) |  |
|  | ≥ 16 | 157 | 84 (54) | 1.26 (0.91 to 1.75) |  | 131 | 68 (52) | 0.93 (0.63 to 1.37) |  |

OR (95% CI): unadjusted odds ratio with 95% confidence intervals,
aOR (95% CI): adjusted odds ratios with 95% confidence intervals,
\* Univariable sample size for outcomes ordered as in table (% of vaccinated cases) = 8824 (36%), 110 (28%), 9952 (41%), 10018 (41%), 3221 (13%), † Multivariable sample size with outcomes ordered as in table (% of vaccinated cases) = 7725 (32%), 100 (25%), 8807 (36%), 8855 (36%), 2856 (12%) ‡ Death due to TB in those who died and where cause of death was known

### Sensitivity analysis of the missing data using multiple imputation

We found that repeating the analysis with an imputed data set had some effect on the results from the complete case analysis. There was a decrease in the accuracy of effect size estimates for BCG vaccination, some increase in p-values (supplementary table S5). However, none of the estimated effects changed their direction, and there were no detectable systematic changes in the results.

For the secondary exposure variables (years since vaccination and age at vaccination, (supplementary table S6 and supplementary table S7), we found a change in direction of the point estimate between years since vaccination and all-cause mortality and recurrent TB, but similar results for age at vaccination and outcomes.

**Supplementary table S5:**
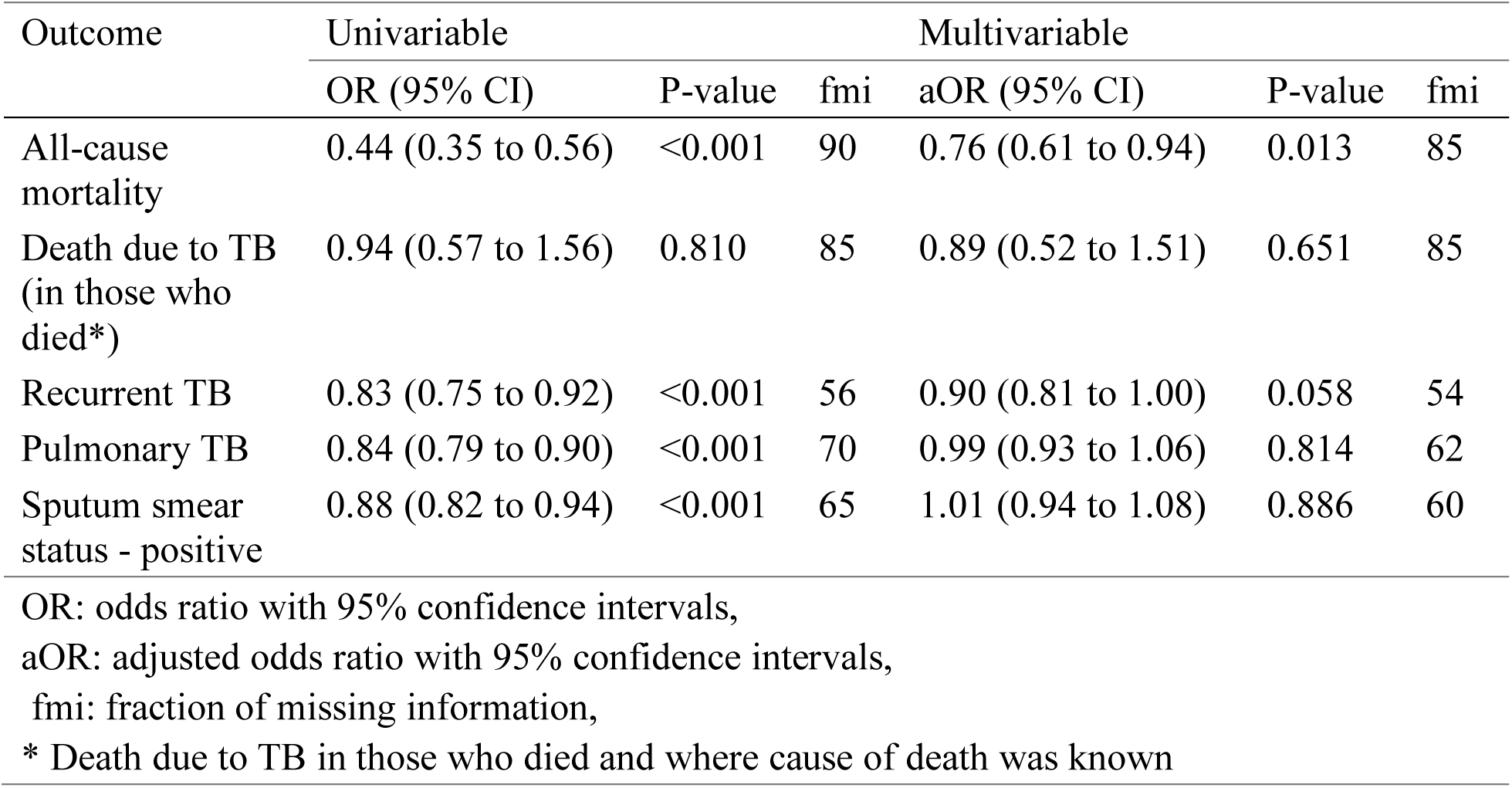
Summary of associations between BCG vaccination and all outcomes, using pooled imputed data.

**Supplementary table S6:** Summary of associations between years since vaccination and all outcomes, using pooled imputed data. There was insufficient data to estimate an effect for deaths due to TB (in those who died)

| Outcome | Univariable |  |  | Multivariable |  |  |
| --- | --- | --- | --- | --- | --- | --- |
|  | OR (95% CI) | P-value | fmi | aOR (95% CI) | P-value | fmi |
| All-cause mortality | 3.28 (1.85 to 5.79) | <0.001 | 50 | 12.19 (3.48 to 42.64) | <0.001 | 70 |
| Death due to TB (in those who died*) | *Insufficient data* | - | 0 | *Insufficient data* | - | 0 |
| Recurrent TB | 1.29 (1.00 to 1.66) | 0.050 | 39 | 0.81 (0.59 to 1.11) | 0.187 | 44 |
| Pulmonary TB | 0.58 (0.52 to 0.66) | <0.001 | 33 | 0.99 (0.84 to 1.17) | 0.913 | 40 |
| Sputum smear status - positive | 0.99 (0.82 to 1.19) | 0.891 | 70 | 0.95 (0.77 to 1.18) | 0.648 | 60 |

OR: odds ratio with 95% confidence intervals,
aOR: adjusted odds ratio with 95% confidence intervals,
fmi: fraction of missing information,
\* Death due to TB in those who died and where cause of death was known

**Supplementary table S7:**
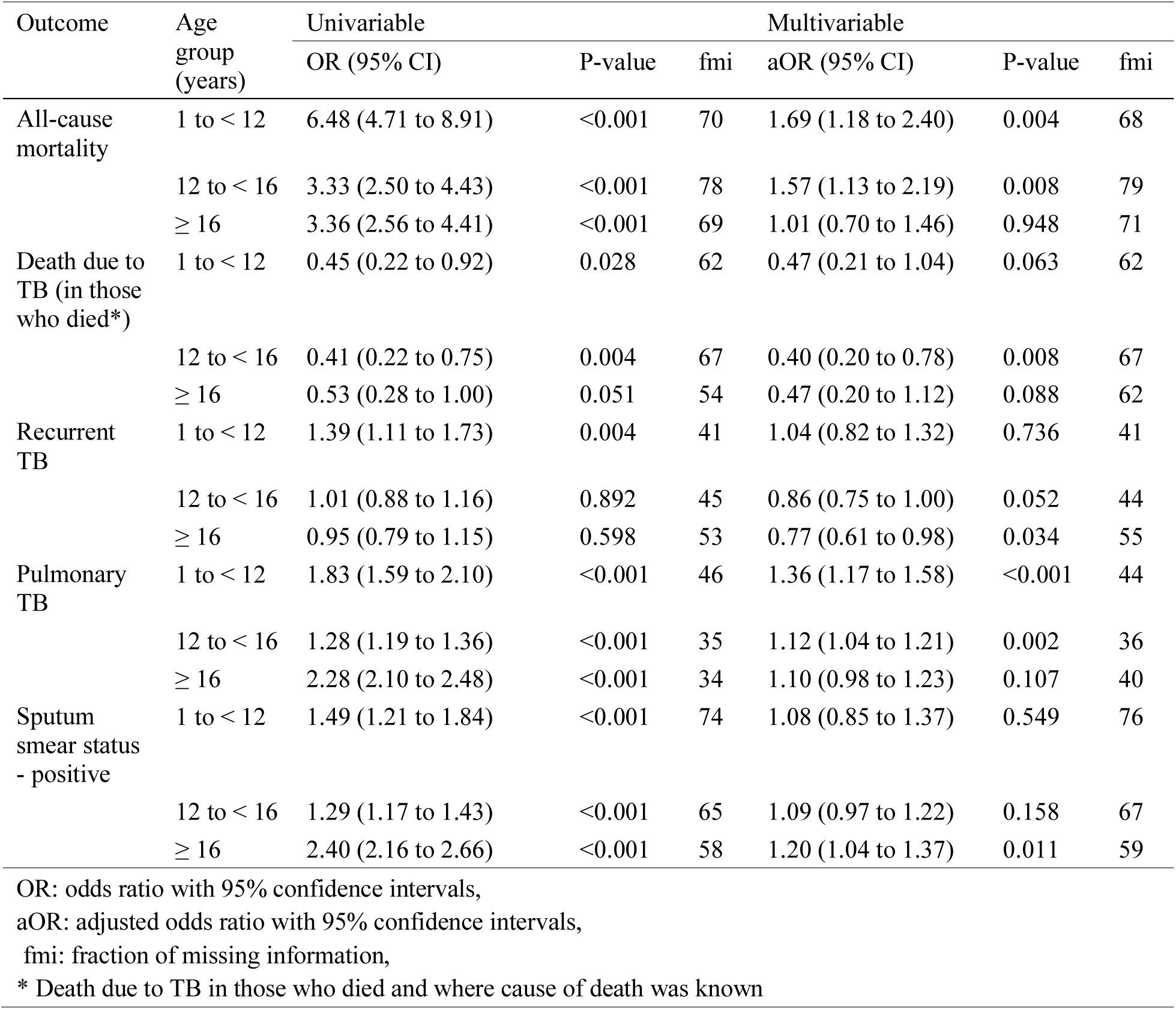
Summary of associations between age at vaccination and all outcomes, using pooled imputed data (reference is vaccination at <1 year).

### Sensitivity analysis of the study population

**Supplementary table S8:**
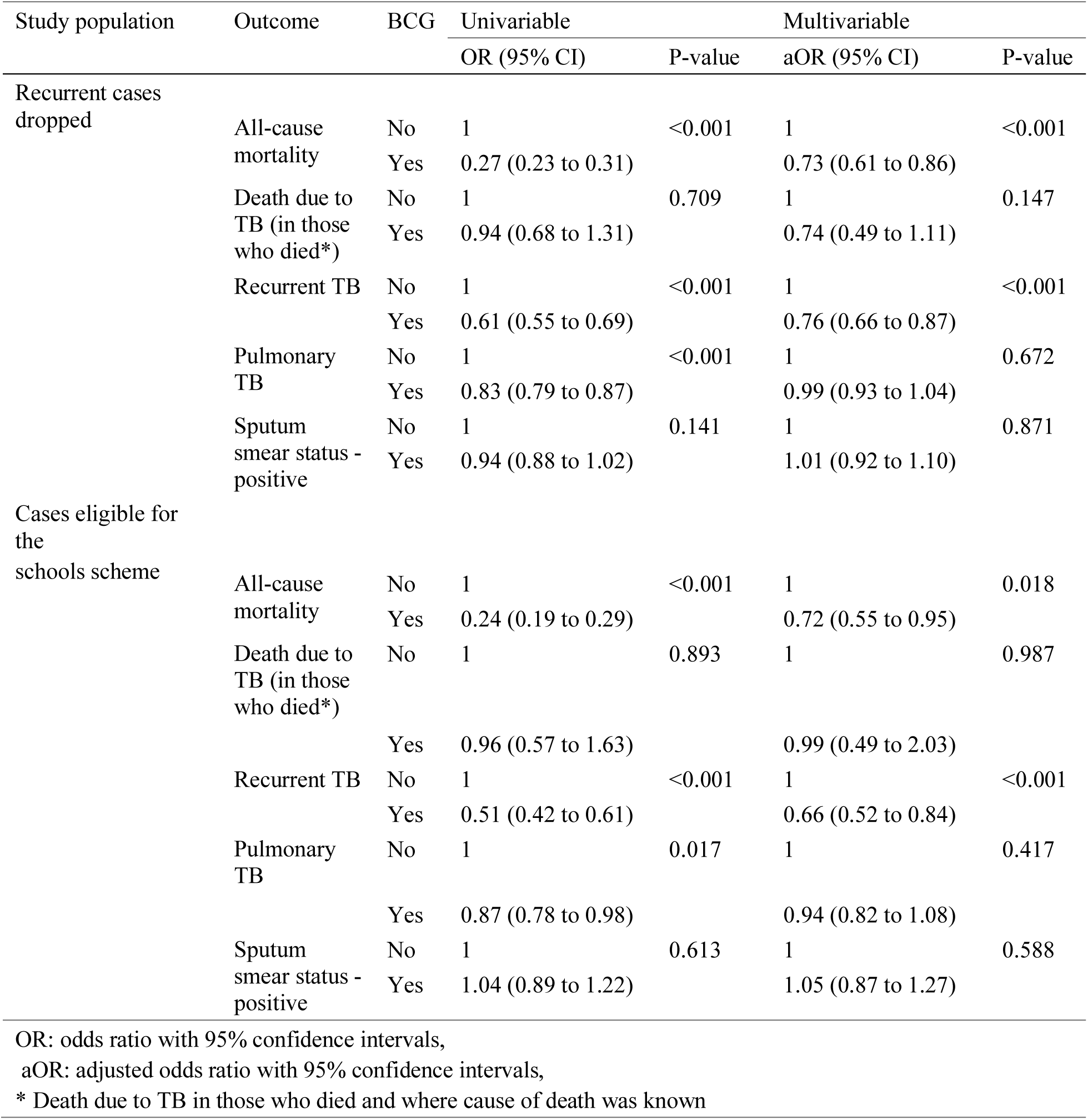
Summary of associations between BCG vaccination and all outcomes; cases that have no recurrent flag in the ETS (50407), and cases that would have been eligible for the BCG schools scheme (9943). Those defined to be eligible for the schools scheme are the UK born, that were aged 14 or over in 2004

### Estimated power

Power estimates were calculated in both the univariable and multivariable datasets for all outcomes. This represents an over estimate of the statistical power, as only a single exposure variable was accounted for, and age at vaccination has been simplified to a binary vaccinated at birth variable (supplementary table S9).

**Supplementary table S9:**
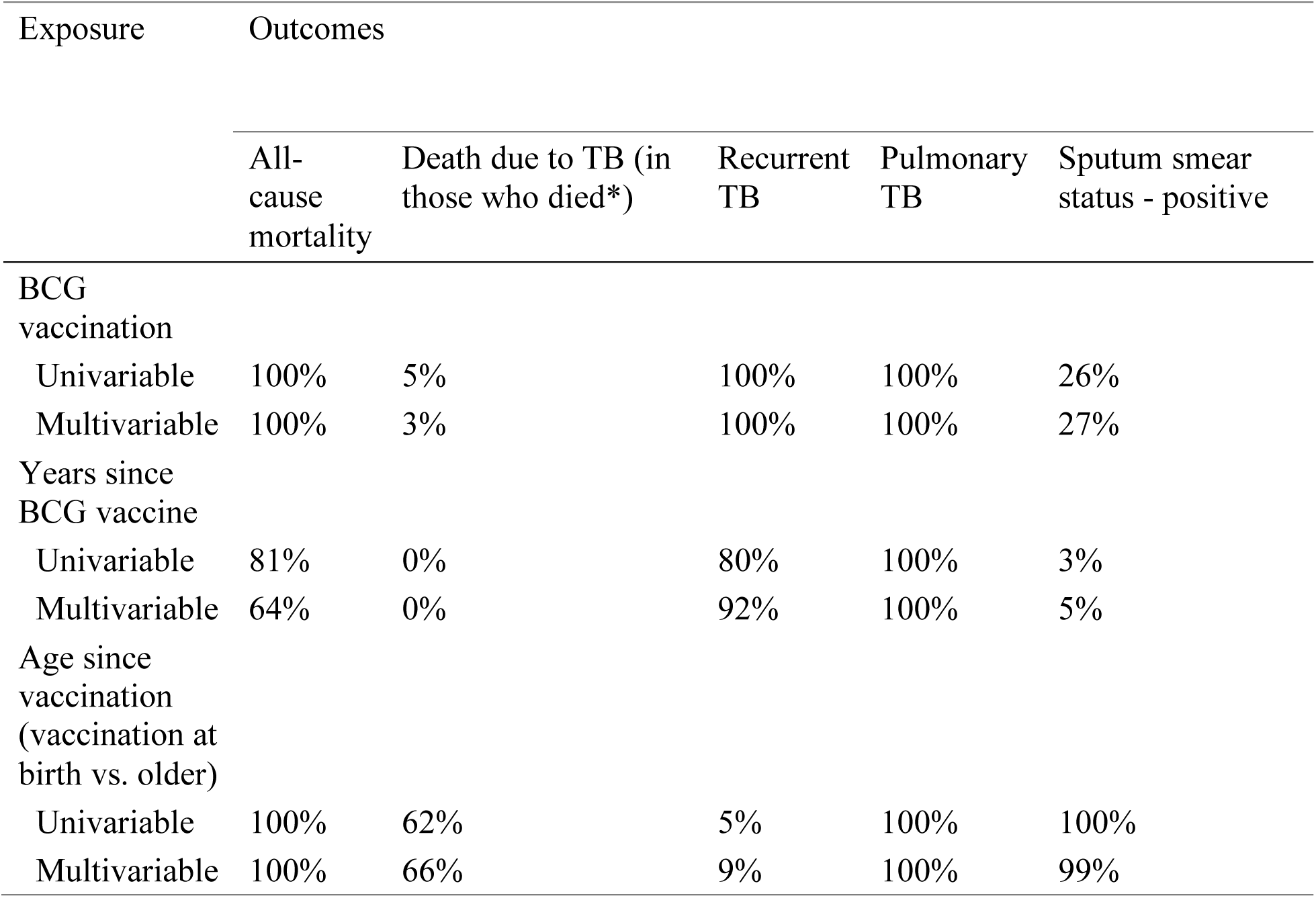
Summary of the estimated power for each analysis, for both the univariable and multivariable data sets, with alpha set as 0.05. Power estimates assume a single exposure variable, and age at vaccination has been simplified into a binary vaccinated at birth variable

| Exposure | Outcomes |  |  |  |  |
| --- | --- | --- | --- | --- | --- |
|  | All-cause mortality | Death due to TB (in those who died*) | Recurrent TB | Pulmonary TB | Sputum smear status - positive |
| BCG vaccination |  |  |  |  |  |
| Univariable | 100% | 5% | 100% | 100% | 26% |
| Multivariable | 100% | 3% | 100% | 100% | 27% |
| Years since BCG vaccine |  |  |  |  |  |
| Univariable | 81% | 0% | 80% | 100% | 3% |
| Multivariable | 64% | 0% | 92% | 100% | 5% |
| Age since vaccination (vaccination at birth vs. older) |  |  |  |  |  |
| Univariable | 100% | 62% | 5% | 100% | 100% |
| Multivariable | 100% | 66% | 9% | 100% | 99% |

